# The CB1 Negative Allosteric Modulator PSNCBAM-1 Reduces Ethanol Self-Administration via a Nonspecific Hypophagic Effect

**DOI:** 10.1101/2023.06.30.547272

**Authors:** Harley M. Buechler, Mousumi Sumi, Indu Mithra Madhuranthakam, Christa Donegan, Frank DiGiorgio, Alisha A Acosta, Sarah Uribe, Mohammad A Rahman, Alison Sorbello, Bradford D. Fischer, Thomas M. Keck

## Abstract

Alcohol use disorder (AUD) affects more than 15 million people in the United States. Current pharmacotherapeutic treatments for AUD are only modestly effective, necessitating the identification of new targets for medications development. The cannabinoid receptor type 1 (CB1) has been a target of interest for the development of medications for substance use disorders and other compulsive disorders. However, CB1 antagonists/inverse agonists (e.g., rimonabant) have severe side effects that limit their clinical utility, including anxiety, depression, and suicide. Recent development of CB1 negative allosteric modulators (NAMs), including PSNCBAM-1, may provide an alternative mechanism of attenuating CB1 signaling with reduced side effects. PSNCBAM-1 has not yet been evaluated for effects in models of AUD. In this study, we investigated the effects of the CB1 NAM, PSNCBAM-1, in rodent models of AUD using adult male mice. PSNCBAM-1 dose-dependently attenuated oral ethanol self-administration (8% w/v ethanol in water), significantly reducing ethanol rewards at a dose of 30 mg/kg, but not at 10 or 18 mg/kg. PSNCBAM-1 also dose-dependently attenuated palatable food self-administration (diluted vanilla Ensure), significantly reducing food rewards at 18 and 30 mg/kg PSNCBAM-1. PSNCBAM-1 did not affect conditioned place preference for 2 g/kg ethanol. These results suggest PSNCBAM-1 reduces ethanol-taking behavior via a nonspecific hypophagic effect and does not reduce the rewarding effects of ethanol.

## Introduction

Alcohol use disorder (AUD) is a serious health condition with massive global impact, characterized by uncontrollable alcohol consumption due to physical and emotional dependence on alcohol and negative effects when alcohol is not consumed.^1^ The 2021 National Survey on Drug Use and Health (NSDUH) determined that 28.6 million adults in the United States aged 18 and above suffered from AUD in the prior year. Globally, in 2019, alcohol consumption was the cause of death for 2.07 million men and 374,000 women.^2^ Current treatments for AUD have low rates of success, indicating a clear need for new pharmacotherapeutics.

The cannabinoid receptor type 1 (CB1), a G protein-coupled receptor (GPCR), expressed in the peripheral and central nervous system, is an intriguing target for development of medications for substance use disorders.^3-4^ The endogenous cannabinoid system signals primarily via CB1 and the cannabinoid receptor type 2 (CB2). CB1 is widely expressed throughout the brain, regulating learning, memory, decision-making, pain, and energy metabolism.^5^ CB1 signaling controls a wide variety of physiological functions, such as food intake, energy balance, cardiovascular functions, reproductive functions, immune modulation, and cell apoptosis.^6^

Neuronal pathways that receive signals through CB1s in the CNS contribute to the rewarding effects of certain non-drug stimuli and many addictive drugs, leading to their excessive intake and pathological consequences. Recent studies have found that CB1 activity can cause a shift from normal behavior to impaired decision making and repetitive drug intake; pharmacologically targeting CB1 is one strategy to reverse these effects and ultimately treat substance use disorders.^4, 7^

Rimonabant (SR1417116A) was the first clinically available, potent, selective, orally active CB1 receptor antagonist.^8^ Both *in vitro* and *in vivo* studies show that rimonabant antagonizes the behavioral and pharmacological effects induced by CB1 agonists.^9-11 12^ An appetite suppressant that was shown to cause weight loss and improve cardiovascular risk factors like dyslipidemia, waist circumference, and blood pressure, rimonabant was initially approved in Europe in 2006 for the treatment of obesity.^13 14^ In 2008, the drug was withdrawn from the market due to serious psychiatric adverse effects, such as suicidal thoughts, anxiety, and depression.^14^

In studies relevant to AUD, rimonabant reduced alcohol-taking and -seeking behaviors.^15-17^ For instance, in studies involving selectively bred Sardinian alcohol-preferring (sP) rats, rimonabant reduced voluntary alcohol consumption.^18-20^ Furthermore, clinical studies of rimonabant revealed its effectiveness in smoking cessation and reducing alcohol dependence.^21-22^ These findings suggest that pharmacotherapies that attenuate CB1 signaling hold promise as potential therapeutic agents in the treatment of AUD.

The clinical failure of rimonabant is typically attributed to its full antagonism, and likely inverse agonism, of CB1 signaling, producing severe and life-threatening mood disorders.^14, 22-23^ To avoid serious adverse events, several research teams have focused on developing novel CB1 negative allosteric modulators (NAMs) with the idea that NAMs would attenuate CB1 signaling in an activity-dependent manner, providing some of the desired therapeutic effects with reduced side-effect profiles.^24-25^

PSNCBAM-1 is classified as a NAM based on its effects in efficacy assays, although it enhances agonist binding.^25-27^ PSNCBAM-1 reduces the overall activation of CB1 by agonists by modulating receptor internalization and thus its effects on cAMP accumulation, overall decreasing the cell’s response to CB1 activation. ^27-28^ Other CB1 NAMs share this set of pharmacological characteristics.^29^ PSNCBAM-1 has not yet been evaluated for effects in rodent models of AUD. This study aimed to investigate the effects of PSNCBAM-1 in models of AUD using adult male mice, with the working hypothesis that PSNCBAM-1 would reduce alcohol reward and alcohol-taking and - seeking behavior. To test this hypothesis, we evaluated the effects of PSNCBAM-1 in ethanol conditioned place preference (CPP) and self-administration studies.

## MATERIALS AND METHODS

### Animals

All studies used drug-naïve male C57BL/6 mice, starting approximately 8 weeks of age. Mice were obtained from Charles River Laboratories (Wilmington, MA). All studies were carried out in the vivarium at Cooper Medical School of Rowan University (CMSRU). Mice were caged in polycarbonate cages (four animals per cage) in a temperature- and humidity-controlled vivarium, with *ad libitum* access to food and water, and enrichment provided by paper Bio-Huts and/or nestlets. All studies were conducted in procedure rooms separate from the housing facility during the light phase of the light/dark cycle (lights on at 7:00, lights off at 19:00). Animals were weighed every day and evaluated for general health and behavioral parameters. All the experiments were conducted in accordance with the Guide for the Care and Use of Laboratory Animals and were approved by the Institutional Animal Care and Use Committee at Rowan University. The CMSRU animal facility of Rowan University is accredited by the Association for Assessment and Accreditation of Laboratory Animal Care International.

### Drugs

PSNCBAM-1 (1-(4-chlorophenyl)-3-(3-(6-(pyrrolidin-1-yl)pyridin-2-yl)phenyl)urea; Tocris) was dissolved in 10% DMSO, 10% Tween 80, and 80% saline vehicle for all studies, using sonication to achieve a clear solution. PSNCBAM-1 was administered via i.p. injections in all experiments at a volume of 10 mL/kg.

190 proof ethanol was purchased from Koptec. For CPP experiments, ethanol was mixed with physiological saline to achieve the desired mass/volume dilution and administered via i.p. injections. For self-administration experiments, ethanol was diluted in Vanilla Ensure™ and water to achieve the desired mass/volume dilution and was administered orally via operant chamber delivery. All dilutions were made weekly and stored at 4 °C between experiments.

### Behavioral Procedures

#### Locomotor Activity

This test was conducted to determine whether PSNCBAM-1 induced substantial locomotor disruptions in mice.

##### Apparatus

Four 40 × 40 cm^2^ modular open field instruments were organized on a vertical shelving unit, two per shelf, for locomotor experiments. A small fluorescent light and a USB camera connected to a PC running Any-maze software were placed above each chamber. Each test apparatus and floor insert were cleaned with 70% isopropyl alcohol and allowed to dry completely before and after all training and testing processes including between each animal in serial testing.

##### Treatment Groups

This study used 32 drug-naïve male C57BL/6 mice. An initial set of 8 vehicle animals were compared to 8 receiving 30 mg/kg PSNCBAM-1. A later study compared 8 vehicle animals to 8 receiving 18 mg/kg PSNCBAM-1. No differences were seen between the vehicle groups so all data were collapsed.

##### Testing

All animals were exposed to the open field twice over the course of two days prior to testing to minimize novelty-induced locomotor activity. Animals were given a saline injection after 15 minutes in the open field during these 55-minute pre-exposures to acquaint them with the handling and injection technique. During these pre-exposures, behavior was not recorded. On the test day, animals were given I.P. injections of 10 mg/kg PSNCBAM-1 or 30 mg/kg PSNCBAM-1 or vehicle mixture and placed in the open field chambers to record their behavior and activity.

##### Analysis

Locomotor activity for each animal was recorded in 5 min intervals. Overall locomotor activity is represented as the total distance traveled following PSCNBAM injection.

#### Conditioned Place Preference (CPP)

CPP is a method for evaluating the subjective effects of drugs of abuse. The CPP test works on the idea that primary reinforcers like legal or illegal drugs, food, water, or sex are coupled with contextual stimuli that gain secondary reinforcing characteristics through Pavlovian contingency.

##### Apparatus

The three-chamber CPP apparatus (Med Associates, Fairfax, VT) placed in sound attenuating cabinets and connected to a computer running MED-PC software (version 5). Each apparatus has two larger compartments with white or black walls and different floor grates. These compartments are connected via a smaller central gray compartment with closable sliding doors which could be used to block or allow the free movement of the mice into and out of the white and black compartments.

##### Initial Preference

Before conditioning/preference training, animals were tested for a predisposed side preference by placing mice in the chambers with full access to all the compartments for 30 min, to remove any bias from the experiment. The ratio of time spent in one test compartment over the total time spent in the central compartment was calculated as a measure of initial compartment preference. Time spent in the central compartment was recorded but not considered in the calculation. Twelve animals with an initial preference of >65% for one test compartment were excluded from training or further testing. The drug-paired and vehicle-paired compartments for each animal were assigned using a Latin square design, which evenly distributed compartment pairing, box placement, and first-day treatment among all treatment groups, as per an unbiased experimental procedure.

An unbiased procedure was used to divide animals into separate treatment groups and plan each training regimen. While biased techniques can provide increased sensitivity for detecting medication effects dependent on the animal’s baseline motivational states, or for detecting anxiolytic and anti-aversive drug effects, they also have a larger false-positive rate, which we wanted to minimize.

##### Treatment Groups

29 mice were ultimately used for this study. Animals were divided into three groups following initial preference testing. The groups received 18 mg/kg PSNCBAM-1 (n=10), 30 mg/kg PSNCBAM-1 (n=10), or vehicle (n=9) 5 min prior to the reinforcer (2.0 g/kg ethanol) for training days on the ethanol-paired side. Animals received vehicle doses prior to training on the saline-paired side.

##### Drug Conditioning

10 training sessions were conducted over a 10-day period for the primary acquisition or conditioning to take place. Animals were given drug or vehicle injections immediately before placing the animal inside the CPP apparatus and confined in the specified test chamber for 30 minutes during training. Drug and vehicle exposures alternated daily, with the pattern of exposures counterbalanced within groups across days and apparatuses.

##### Preference Test

In a CPP expression trial experiment, trained mice were given no injection and were given free access to the complete apparatus (including both drug- and vehicle-paired compartments) for 30 minutes.

##### Analysis

For both the initial (Δ_pretest_) and final preference (Δ_posttest_) tests, a preference score was generated by subtracting the seconds spent in the vehicle-paired chamber from the seconds spent in the drug-paired compartment (time in the central compartment was ignored). The following formula was used to compute an overall preference score for a specific pharmacological dosage: Preference score (sec) = Δ_postest_ − Δ_pretest_

#### Food and Ethanol Self-Administration

Self-administration studies were conducted to examine the effects of PSNCBAM-1 on operant behavior reinforced by a palatable food reward or ethanol. properties of neuropharmacological drugs/compounds. Animals were trained to nose poke to obtain food or ethanol. After sufficient training, animals were injected with the drug under investigation and evaluated for its anti-addiction property in terms of reduced ethanol self-administration.

##### Apparatus

Self-administration training and testing used mouse operant chambers (Med Associates, Fairfax, VT) placed in sound-attenuating cabinets and connected to a computer running MED-PC software (version 4). Each operant chamber featured two nose poke holes equipped with infrared beams. A liquid dipper was located between the two nose poke holes, connected via tubing to a pump and syringe that delivered the programmed reward. For both food and ethanol self-administration studies, the left nose poke hole was paired with food or ethanol reinforcers, while the right nose poke hole had no programmed consequences. Upon earning a programmed reward, the syringe pump discharged the liquid reinforcer for 3 seconds (delivering an approximate 0.1 mL volume). All rewards were accompanied by light and tone stimuli during the duration of the 3 second reward delivery. There was no programmed timeout.

##### Treatment Groups

Two separate groups of 8 mice were used for food and ethanol self-administration studies, respectively. Animals in the ethanol group received identical food self-administration training before a fading procedure was used to transition them to ethanol.

##### Food Restriction

Before each training or testing session, mice were food restricted for approximately 21 hours, but had *ad lib* access to water. Every day, mice were weighed and held within 10% of their free-feeding weight. Animals were allowed *ad lib* access to chow for 1 hour after each training or testing session. In prior pilot studies, we were unsuccessful in achieving reliable ethanol self-administration in mice without maintaining food self-administration.

##### Training

All mice were trained initially to nose poke for a palatable food reward, diluted vanilla Ensure™ (50:50 water:Ensure), initially using a fixed-ratio 1 (FR1) in which a single nose poke into the reinforcer-paired nose poke hole resulted in one reward delivery. As each mouse earned >90 of 100 possible food rewards in a 2-hour period over three days on a given FR level, the FR was successively increased, from FR2 to FR3 and FR4. Once stable FR4 food self-administration was achieved, animals moved onto drug testing (food self-administration group) or an ethanol fading procedure.

##### Ethanol Fading Procedure

Ethanol was gradually introduced into the diluted vanilla Ensure™, incrementally increasing the percentage of ethanol from 2%, to 4%, to 6%, hen to 8% w/v (equivalent to 10.13% v/v), replacing the water in the baseline 50:50 water:Ensure mixture. Then, the Ensure was incrementally replaced with water until reaching a final 8% w/v mixture of ethanol and water in each earned reward. Animals progressed to the next dilution after earning a consistent number of earned reinforcers (± 10%) for at least 3 straight days. Baseline 8% w/v ethanol self-administration averaged ∼10 rewards over a 2-hour training session; mice appeared visibly intoxicated at the end of training sessions (altered gait, sluggish responses) due to the relatively high training dose

##### Testing

After meeting FR4 training criteria for food or ethanol intake, mice were tested with varying doses of PSNCBAM-1 (10 mg/kg, 18 mg/kg, 30 mg/kg, or vehicle) using a Latin square design. PSNCBAM-1 was given i.p. 5 minutes before placing the animal inside the operant chamber and beginning a standard 2-hour procedure.

#### Data Analysis

Data from all studies was collected and analyzed (GraphPad Prism 6) using one-way or repeated-measures ANOVA, as appropriate, followed by pre-planned Bonferroni analyses.

## Results

The effects of PSNCBAM-1 were tested in initial control studies to determine whether PSNCBAM-1 would produce general behavioral disruptions that might complicate interpretations of other planned behavioral tests. Initial control studies indicated that 10 and 30 mg/kg, i.p., PSNCBAM-1 did not significantly disrupt locomotor activity in the open field (**Figure 1**). One-way ANOVA analysis of total post-injection locomotor activity revealed no significant effect of treatment (F(2,29) = 0.5594, p > 0.5).

**Figure 1.**
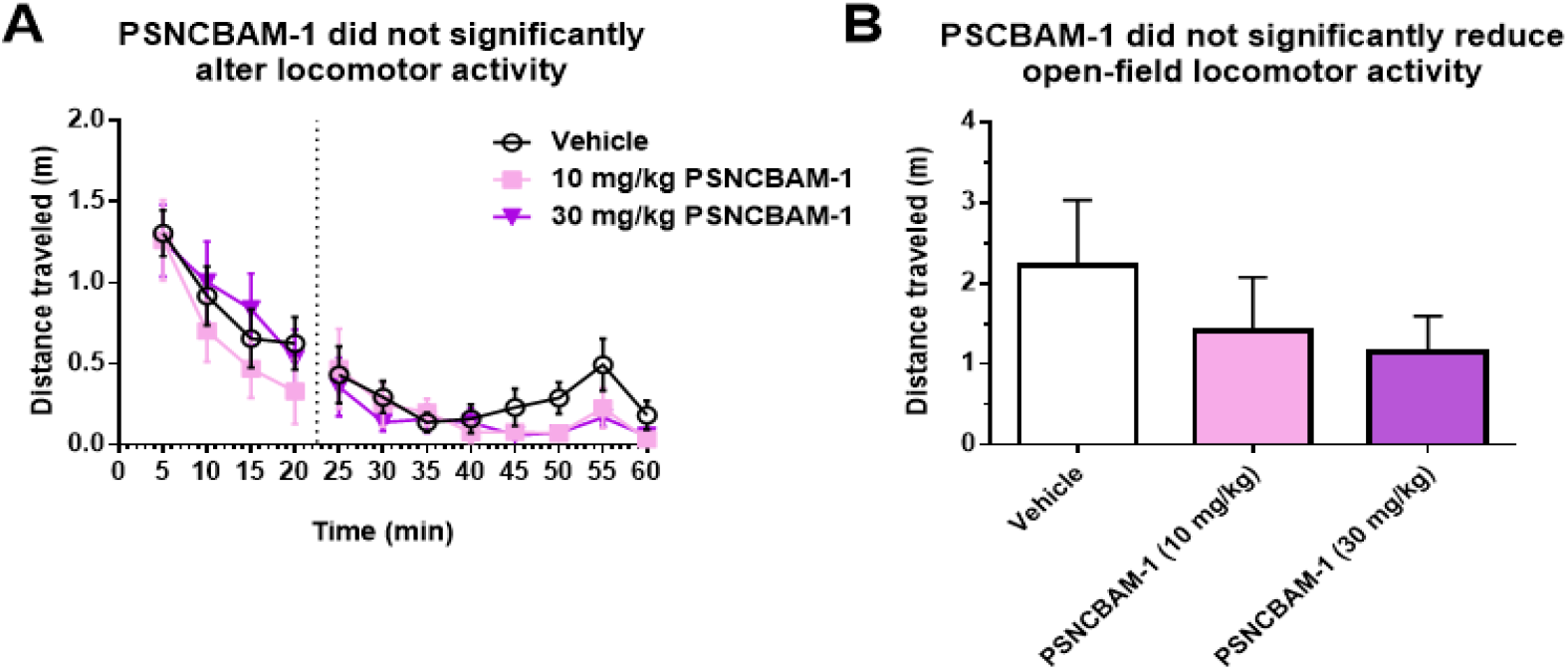
PSNCBAM-1 did not significantly alter locomotor activity in the open field. (A) 20 minutes after introduction into the open field, mice were injected with 10 or 30 mg/kg PSNCBAM-1 or vehicle and locomotor activity was recorded for an additional 40 minutes. (B) Overall post-injection distance traveled was not significantly different across treatments. All data are presented as means ± SEM.

**CPP:** Three different dosages, 10 mg/kg, 18 mg/kg, and 30 mg/kg, in addition to a vehicle mixture, were examined as a pretreatment to a 2.0 g/kg dose of ethanol or saline vehicle for the PSNCBAM-1 CPP tests (**Figure 2**). PSNCBAM-1 administration during conditioned place preference training had no effect on the rewarding value of 2.0 g/kg ethanol. A one-way ANOVA study of preference for the ethanol-paired compartment indicated that PSNCBAM-1 administration had no significant impact (F (2,26) = 0.3469, p > 0.7).

**Figure 2.**
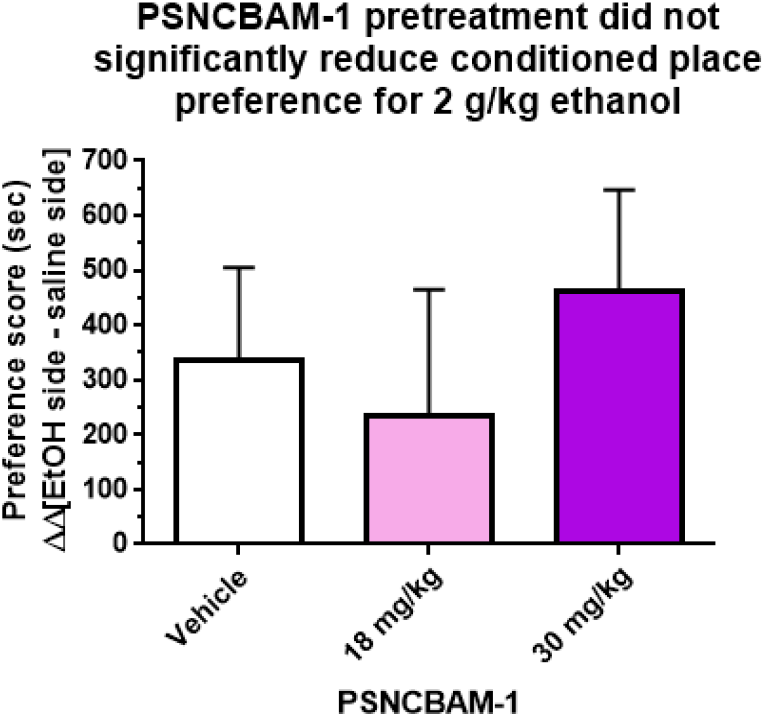
PSNCBAM-1 did not significantly alter conditioned place preference for 2 g/kg ethanol. Pretreatment with 18 or 30 mg/kg PSNCBAM-1 prior to ethanol treatment did not attenuate acquisition of ethanol place preference compared to vehicle control. All data are presented as means ± SEM.

### Self-administration

All mice were trained initially to self-administer a 50% Ensure/50% water combination for food self-administration. The mice were evaluated in a Latin square design with two different dosages of PSNCBAM-1, 10 mg/kg and 30 mg/kg, and vehicle. An intermediate dosage of 18 mg/kg was examined in the second phase of testing. Mice were given i.p. injections of PSNCBAM-1 or vehicle prior to a 2-hour self-administration session on test days. PSNCBAM-1 inhibited palatable food self-administration in a dose-dependent manner, considerably lowering food rewards at 18 mg/kg and 30 mg/kg (**Figure 3**). One-way repeated-measures ANOVA revealed a significant effect of PSNCBAM-1 treatment (F (3,18) = 4.264, p = 0.0193). Bonferroni testing indicated a significant difference between vehicle and 30 mg/kg PSNCBAM-1 treatment (t = 3.016, p 0.05).

**Figure 3.**
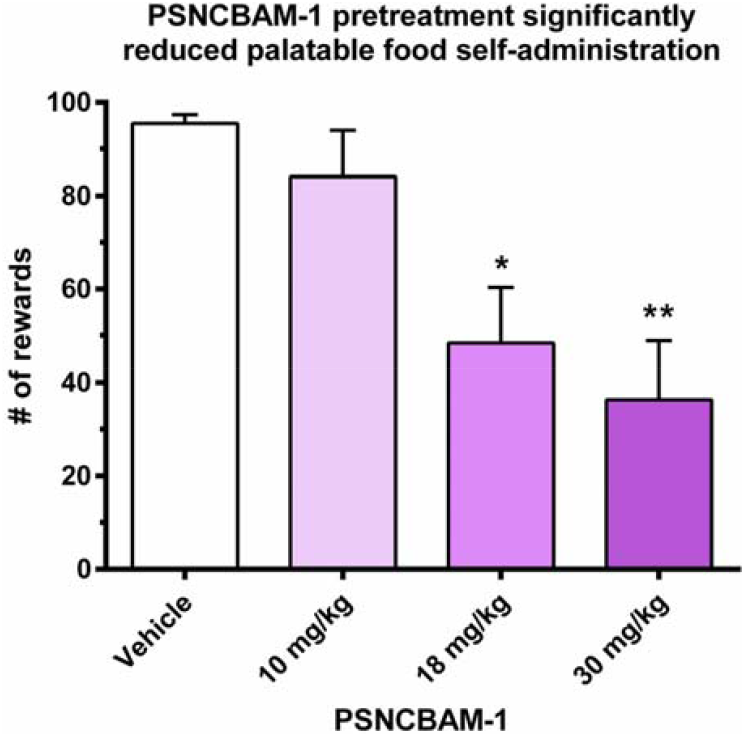
18 and 30 mg/kg, but not 10 mg/kg, PSNCBAM-1 significantly decreased self-administration of a palatable food reward. Pretreatment with 18 or 30 mg/kg PSNCBAM-1 prior to a food self-administration session reduced the number of palatable food rewards received compared to vehicle control. All data are presented as means ± SEM. * p< 0.05, ** p< 0.01, compared to vehicle control.

To determine whether PSNCBAM-1 could reduce ethanol self-administration behavior, mice trained to self-administer 8% (w/v) ethanol were tested with various doses of PSNCBAM-1 in a Latin square design (**Figure 4**). On test days, mice were given i.p. injections of PSNCBAM-1 or vehicle immediately prior to a 2-hour self-administration session. One-way repeated-measures ANOVA revealed a significant effect of PSNCBAM-1 treatment (F(3,31) = 8.410, p = 0.0007). Pre-planned Bonferroni tests revealed a significant difference between vehicle treatment and 18 mg/kg (t = 3.426, p < 0.05) and 30 mg/kg PSNCBAM-1 (t = 4.298, p < 0.01).

**Figure 4.**
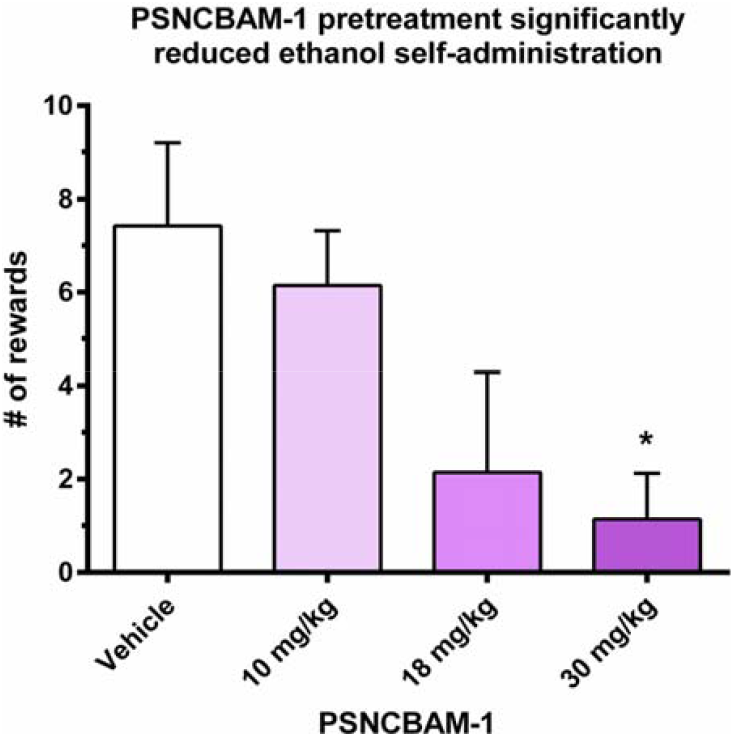
30 mg/kg, but not 10 or 18 mg/kg, PSNCBAM-1 significantly decreased ethanol self-administration. Pretreatment with 30 mg/kg PSNCBAM-1 prior to an ethanol self-administration session reduced the number of ethanol rewards received compared to vehicle control. All data are presented as means ± SEM. * p< 0.05, compared to vehicle control.

## Discussion

Studies have consistently shown that attenuation of CB1 signaling can suppress appetitive behaviors for food as well as drugs of abuse.^4, 30-36^ PSNCBAM-1 is a CB1 NAM that acts centrally and peripherally. In the CNS, PSNCBAM-1 exerts effects opposite to that of CB1 agonists like THC and CP-55,940. Prior to this study, PSNCBAM-1 had never been evaluated for effects in rodent models of AUD.

In these studies, PSNCBAM-1 did not significantly affect the acquisition of place preference to 2.0 g/kg ethanol, suggesting that the doses tested did not alter the rewarding value of 2 g/kg ethanol. In operant studies, PSNCBAM-1 attenuated food and ethanol (8% w/v) self-administration at doses that did not affect CPP. Control experiments indicate that PSNCBAM-1 did not significantly affect locomotor activity in an open-field at doses that altered self-administration behavior. Taken together, these results suggest that PSNCBAM-1-mediated attenuation of ethanol self-administration was not driven by any reduction in the rewarding value of alcohol but rather a general hypophagic response. The results of this study are fully consistent with a prior investigation of PSNCBAM-1 that reported hypophagic effects of PSNCBAM-1 in rodents, reducing food intake and body weight.^26^ Overall, these results do not support the utility of PSNCBAM-1 for AUD treatment.

Recently, a new CB1 NAM (RTICBM-74^37^) was reported to reduce alcohol intake in rats trained to self-administer ethanol.^38^ This point of contrast highlights some limitations to take into consideration when evaluating the results presented here. Importantly, this study only evaluated male mice of a single strain, and all tests were performed within the light part of the light:dark cycle. Male and female mice have different sensitivities to the locomotor effects induced by ethanol^39-40^ and patterns of ethanol self-administration can vary between male and female mice.^41^ Rats and mice also have different capacities for operant learning. Finally, we only tested a single dose of ethanol in our CPP studies—a dose that is at the peak of our dose-response curve in pilot studies—and it is possible that PSNCBAM-1 may have influenced the rewarding value of smaller or larger ethanol doses.

In conclusion, we examined the effects of the CB1 NAM PSNCBAM-1 in mouse models relevant to AUD. Our results showed that PSNCBAM-1 pretreatment did not significantly disrupt locomotor activity or affect the rewarding value of 2.0 g/kg ethanol. PSNCBAM-1 dose-dependently attenuated ethanol self-administration, reducing ethanol rewards at the 30 mg/kg dose, but also dose-dependently attenuated palatable food self-administration, significantly reducing food rewards at the 18 mg/kg and 30 mg/kg doses. These results are most parsimoniously explained by a PSNCBAM-1-mediated effect on general appetitive behavior and not on the rewarding value of ethanol.

## Acknowledgments

This work was supported by internal funds from Rowan University and Cooper Medical School of Rowan University. Equipment used for these studies were purchased with support from DA041560.

## Conflicts of Interest

On behalf of all authors, the corresponding author states that there is no conflict of interest.

## Notes

### Competing Interest Statement

The authors have declared no competing interest.

### Summary of Updates

Minor fixes to clarify methods and clean up typos

